# Comprehensive Fitness Landscape of a Multi-Geometry Protein Capsid Informs Machine Learning Models of Assembly

**DOI:** 10.1101/2021.12.21.473721

**Authors:** Daniel D. Brauer, Celine B. Santiago, Zoe N. Merz, Esther McCarthy, Danielle Tullman-Ercek, Matthew B. Francis

**Affiliations:** Department of Chemistry, University of California Berkeley, Berkeley, CA; Department of Chemical and Biological Engineering, Northwestern University, Evanston, IL; Department of Chemistry, University of California, Berkeley Berkeley, CA; Materials Sciences Division, Lawrence Berkeley National Laboratories, Berkeley, CA

## Abstract

Virus-like particles (VLPs) are non-infections viral-derived nanomaterials poised for biotechnological applications due to their well-defined, modular self-assembling architecture. Although progress has been made in understanding the complex effects that mutations may have on VLPs, nuanced understanding of the influence particle mutability has on quaternary structure has yet to be achieved. Here, we generate and compare the apparent fitness landscapes of two capsid geometries (T=3 and T=1 icosahedral) of the bacteriophage MS2 VLP. We find significant shifts in mutability at the symmetry interfaces of the T=1 capsid when compared to the wildtype T=3 assembly. Furthermore, we use the generated landscapes to benchmark the performance of *in silico* mutational scanning tools in capturing the effect of missense mutation on complex particle assembly. Finding that predicted stability effects correlated relatively poorly with assembly phenotype, we used a combination of *de novo* features in tandem with *in silico* results to train machine learning algorithms for the classification of variant effects on assembly. Our findings not only reveal ways that assembly geometry affects the mutable landscape of a self-assembled particle, but also establish a template for the generation of predictive mutational models of self-assembled capsids using minimal empirical training data.

## Introduction

Virus-like particles (VLPs) have emerged as promising scaffolds for many applications in biotechnology, including vaccine development,^1-3^ targeted drug delivery,^4,5^ and nanoreactor production.^6^ These well-defined, highly symmetrical closed-shell structures self-assemble from discrete protein building blocks that can be engineered to tune the physical and chemical features of the assembly.^7^ While great strides have been made in *de novo* design of self-assembling VLP mimetics,^8–13^ predicting the effects of mutations to the subunit building blocks of a nanocage remains challenging because even small changes can disrupt or drastically alter the complex network of interactions that drive assembly.^14–16^

Systematic study of the effects of mutations on a given protein function has recently been made possible by the advent of deep-mutational scanning, a method that uses next-generation sequencing to assess > 10^5^ protein variants in a single experiment.^17,18^ This technique has previously been applied to self-assembling viral structures such as AAV, HIV,^19,20^ influenza,^21^ and polio^22^ in order to generate fitness landscapes that describe the effects of mutations on viral infectivity. In order to probe the impact of mutations on assembly interactions more directly, we recently developed a strategy called SyMAPS (Systematic Mutagenesis and Assembled Particle Selection), which employs capsid self-assembly as a fitness readout for deep mutational scanning. ^23^ This method enables a quantitative, systematic understanding of how mutations to the subunits of a closed-shell particle affect the final assembly state without relying on a selection that bundles many fitness criteria (e.g. infectivity requires successful viral attachment, replication, and assembly).^24^ SyMAPS has been successfully utilized to quantify the effects of single mutants, epistatic interactions, loop insertions, and peptide extensions in bacteriophage MS2 VLPs, enabling tuned particle thermostability, acid lability, and chemical reactivity.^25–27^

Here, we employ the SyMAPS approach to generate the apparent fitness landscape of a non-native assembly geometry of an icosahedral VLP. Prior work uncovered a point mutant (MS2[S37P]) in the coat protein of the well-studied MS2 bacteriophage VLP that results in a shift in quaternary assembly of the particle.^16^ This shift from T=3 to T=1 icosahedra dramatically reorganizes the capsid assembly while maintaining nearly identical primary, secondary, and tertiary structures of the constituent coat protein subunits (Figure 1).

**Figure 1:**
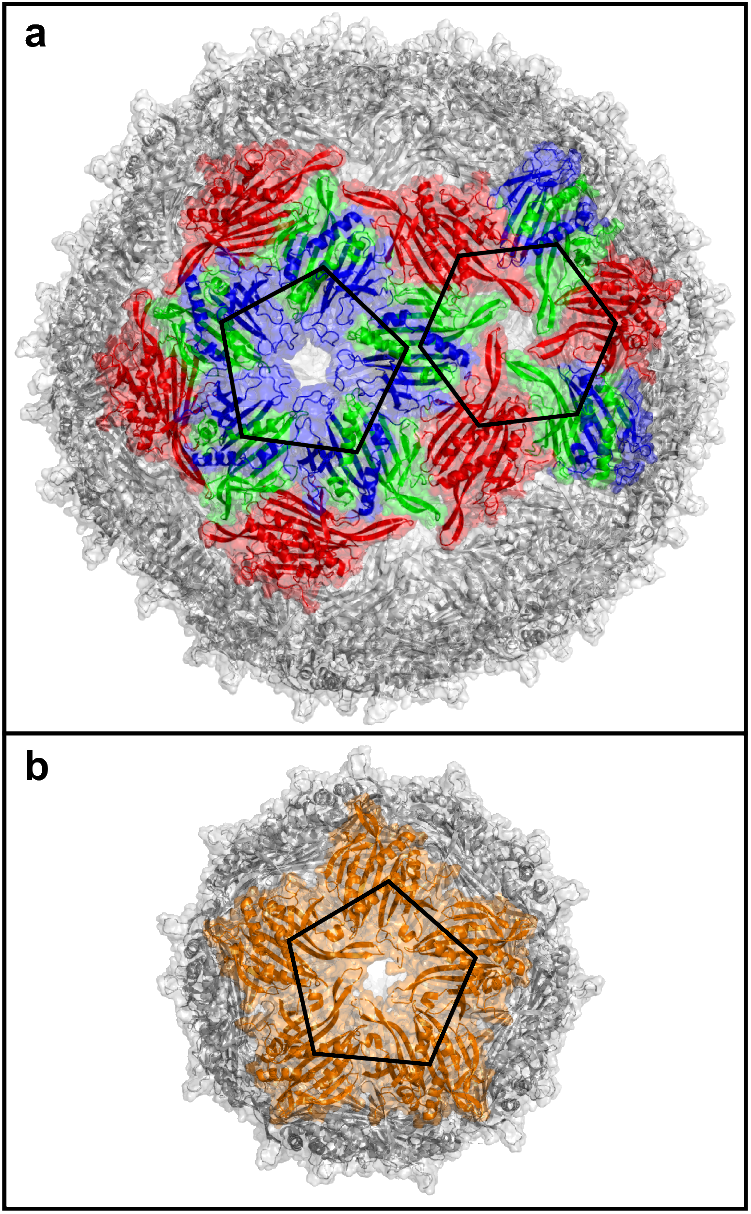
Bacteriophage MS2 virus-like particle assemblies. (a) MS2[WT] T=3 icosahedral capsid is shown (PDB ID: 2MS2). C/C and A/B dimer conformers are shown in red and green/blue, respectively. (b) MS2[S37P] T=1 icosahedral capsid (PDB ID: 4ZOR). All dimers in this capsid geometry are equivalent. Assembly capsomers are outlined in black to show constituent symmetry interfaces of each particle geometry.

By generating a 1-dimensional AFL of MS2[S37P] we were able to uncover unprecedented insight into the underlying principles of assembled structure mutability — we find that core residues and residues at the dimer interface of both VLP geometries share similar mutabilities on average. On the other hand, residues mediating interface between dimer subunits or at multiple contact interfaces are significantly more mutable in the T=1 variant. Interestingly, this holds true for residues both at the quasi-6-fold symmetry interface, which only occurs in the wildtype VLP, and the 5-fold symmetry interface, which is present in both capsid phenotypes. We then use the comprehensive variant data to assess computational methods for determining changes to the folding free energy of a protein when point mutants are introduced, and find that correlation between the predicted folding ΔΔG and the experimental effect on assembly viability is relatively low. Lastly, we combine computational prediction results with experimental fitness landscape results to train machine learning models of assembly state classification (Figure 2). We find that minimal computational and experimental input may be used to generate well-performing classifiers for mutational effects on VLP assembly state.

**Figure 2:**
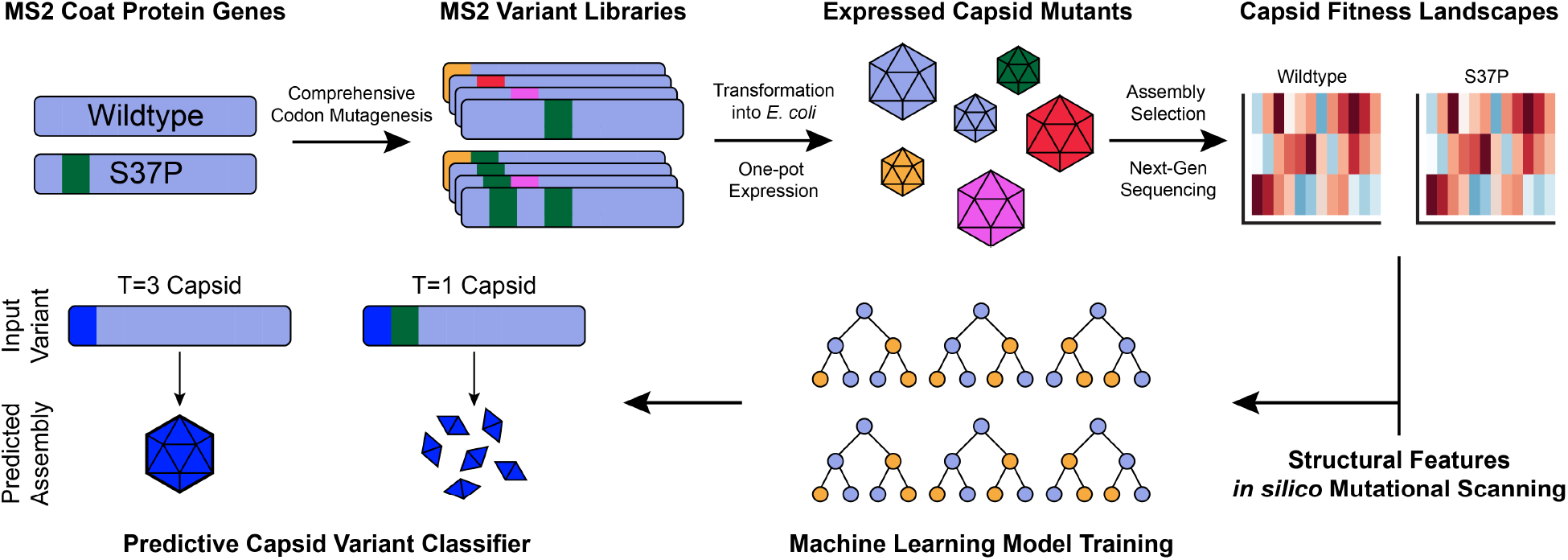
Workflow for generation of capsid apparent fitness landscapes and subsequent assembly classifier generation.

## Results

### Deep Mutational Scan of a Non-Native Virus-like Particle Assembly

In order to explore the effect of quaternary structure on the mutable landscape of self-assembling VLPs, we generated a deep mutational scanning library of the MS2 coat protein with a fixed S37P backbone point mutation. This backbone, MS2[S37P], shifts the MS2 VLP assembly from a 27 nm, T=3 icosahedral geometry to a 17 nm, T=1 icosahedral assembly. ^16^ The SyMAPS platform was used to construct triplicate libraries of the MS2[S37P], as well as the wildtype (MS2[WT]) backbone. Next-generation sequencing of the mutagenized plasmid libraries resulted in excellent coverage of the mutational space, with 94% and 96% of possible variants accounted for in the MS2[S37P] and MS2[WT] libraries respectively. In both cases the majority of variants missing from the library were at the first and last position of the coat protein, which is the result of the first 5 base pairs of the Illumina PE300 MiSeq reads falling below our quality score threshold. The mutagenized libraries were expressed and subjected to size-exclusion chromatography to distinguish between assembling T=1 capsids, assembling T=3 capsids, and non-assembling variants. Assembled capsids encapsidate their encoding mRNA, thus the genotype of each well-formed particle was retained for sequencing. A high-temperature challenge was also introduced to remove capsids with compromised thermostability. Following coat protein expression and capsid enrichment, 72% of MS2[S37P] variants and 84% of MS2[WT] variants were recovered via sequencing and assigned an Apparent Fitness Score (AFS) derived from the log_10_ change in DNA read abundance between the input plasmid library and assembled capsid library. Correlation of AFS between replicates of the MS2[WT] was relatively high (*r*^2^ = 0.70 – 0.83), indicating strong reproducibility and minimal distortions from biological noise (Figure 3a). Interreplicate correlation in the MS2[S37P] libraries was somewhat lower (*r*^2^ = 0.32 – 0.56), possibly due to a lower signal-to-noise ratio, as T=1 capsids encapsidate less genetic material than the wildtype capsid (Figure 3b).^28^ There was a low degree of correlation between the MS2[WT] and MS2[S37P] libraries, meaning that there are both shared mutability preferences and divergences in mutability between the two capsid geometries. A notable difference in AFS distribution of the libraries is derived from their tolerance to deleterious mutations. A large portion of poorly performing variants in the MS2[WT] library (*AFS* = –2.0 – 0.0) drop to the lowest possible fitness score in the MS2[S37P] library (AFS = −4) (Figure 3c). This suggests that the MS2[WT] capsid may retain some level of assembly competency in response to deleterious mutations, while the MS2[S37P] capsid assembly is completely ablated in such cases. This may however also result from lower mRNA packaging per T=1 capsid, thus dropping deleterious mutations below the detection limit.

**Figure 3:**
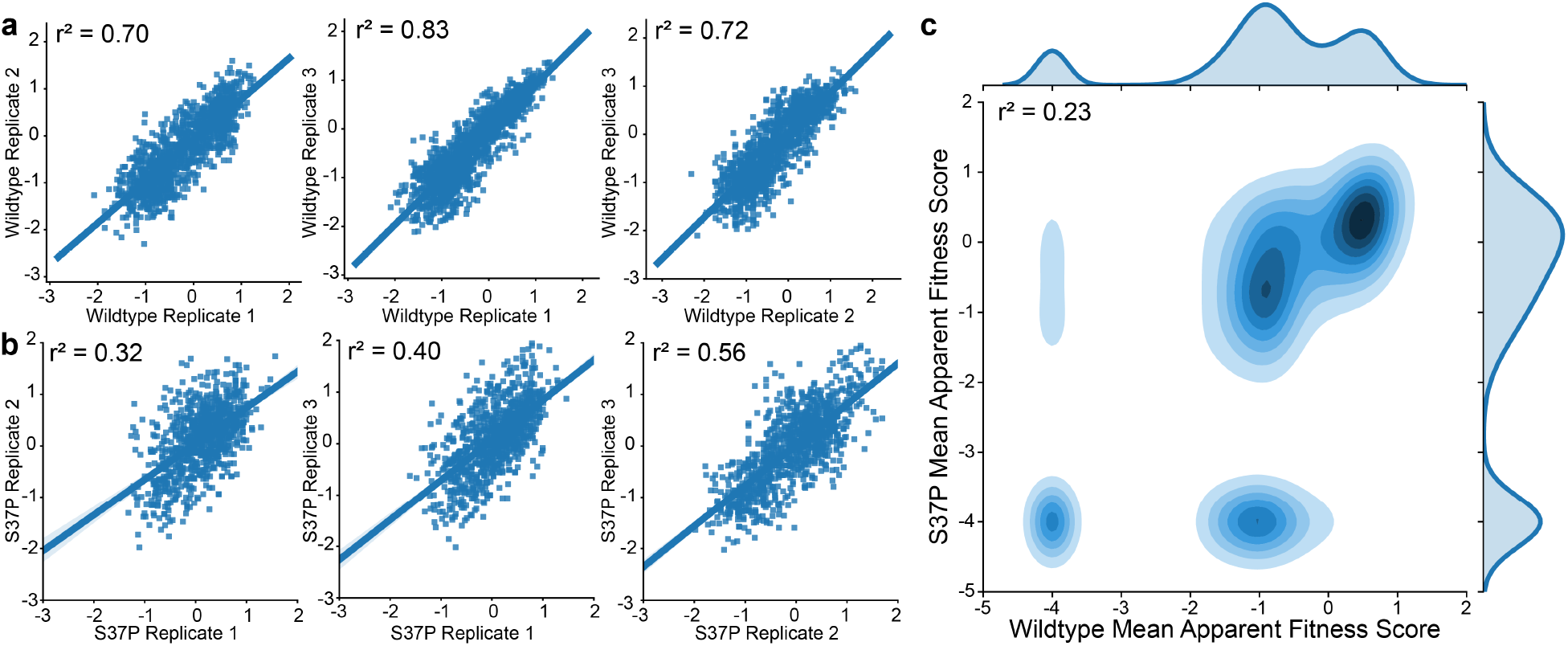
Capsid apparent fitness landscape replicate correlation. Correlations between the three biological replicates of the (a) MS2[WT] and (b) MS2[S37P] libraries are shown. (c) Correlation between the mean apparent fitness scores of each variant in the MS2[S37P] and MS2[WT] libraries. Kernel density estimations of apparent fitness score distributions are shown outside of the axes.

To investigate how the mutational landscape of the T=1 assembly phenotype differs from the wildtype T=3 icosahedron further, Shannon Entropy calculations were performed for each position of the capsid monomer backbone.^29^ This measure of diversity at a given residue has previously been used to generate a Mutability Index (MI) of each backbone position,^23^ which may be used to determine engineering “hotspots” in the capsid coat protein as well as conserved residues likely to mediate key interactions for successful protein folding and particle self-assembly. The difference in MI between the MS2[WT] and MS2[S37P] capsid monomers is shown in (Figure 4a). The majority of the coat protein backbone shows a relatively low difference in MI between the wildtype and S37P libraries, indicating that many of the essential folding and assembly interactions are preserved. Regions of defined secondary structure elements, such as the central portion of the alpha helix spanning residues 105-111, show low mutability in both the T=1 and T=3 capsids. Likewise, the well-studied FG loop region^30,31^ (residues 71-76) of the coat protein retains high mutability in both capsid assemblies. However, individual positions with a significant shift in MI (< −0.2 or > 0.2) span the length of the backbone. The difference in mean fitness at each backbone is shown in (Figure 4b-c). Regions of where the mean AFS of MS2[WT] is notably higher than MS2[S37P] are prevalent, while positions with a higher mean AFS in MS2[S37P] are relatively rare. Mapping of MI to the capsid structures reveals the variety of secondary structural motifs with shifts in mutability between the two capsid phenotypes — residues in beta sheets, alpha helices, turns, and disordered loops show unique preferences in the MS2[S37P] capsid (Figure 4d).

**Figure 4:**
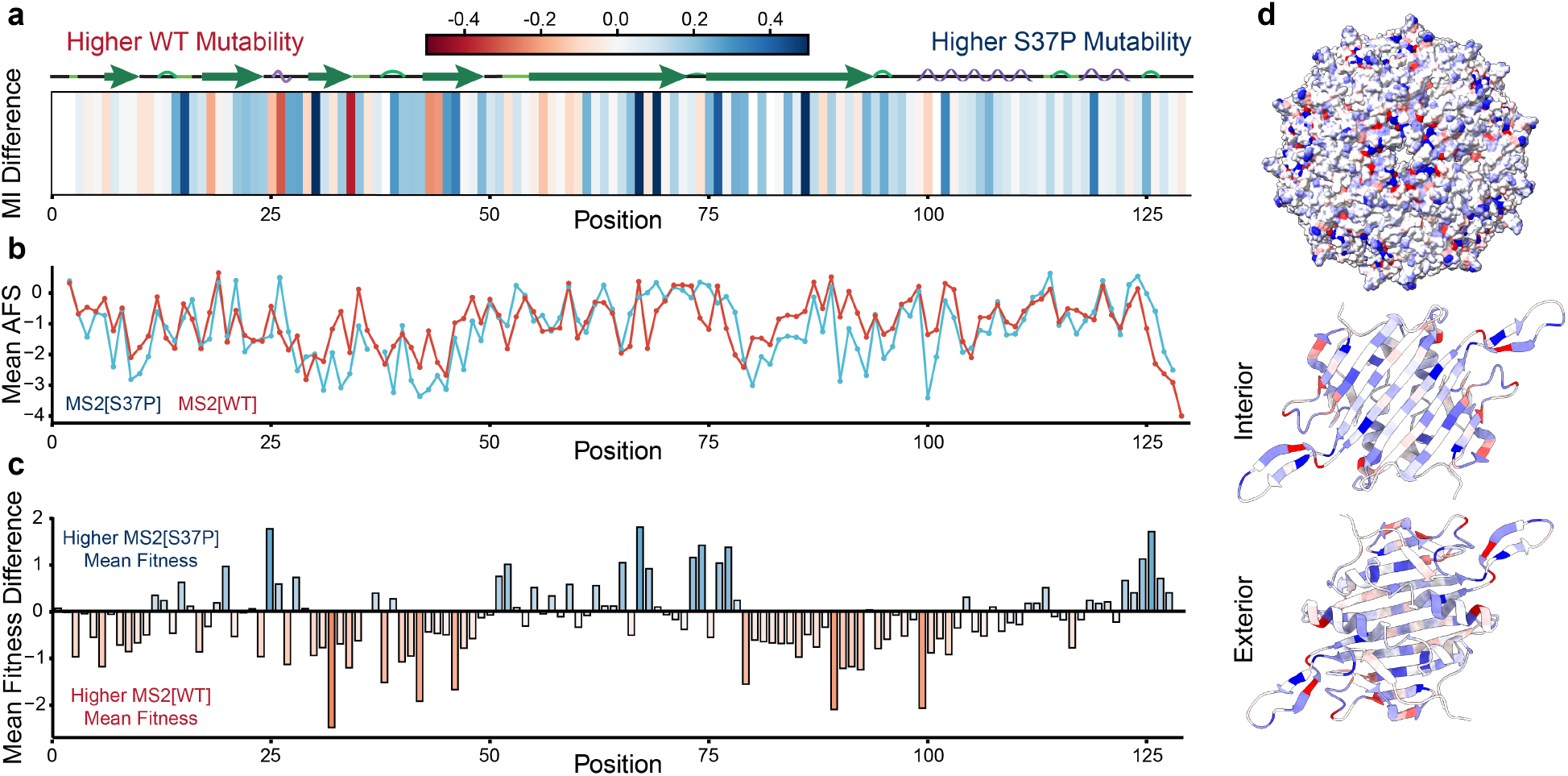
Global mutability trends in MS2 capsid assemblies. (a) Per-residue differences in the mutability index between MS2[WT] and MS2[S37P] capsid backbones. (b) Mean apparent fitness scores (AFS) of all mutations at a given backbone position for both mutagenic libraries. (c) Differences in mean AFS for both capsid libraries. (d) Differrences in mutability index mapped to the MS2[S37P] capsid (PDB ID: 4ZOR) as well as its constituent dimer using ChimeraX. ^32^ In all cases blue represents a higher value in MS2[S37P] and red represents a higher value in MS2[WT].

Surprisingly, it is much more common that positions become more permissive to mutations in the T=1 capsid than become more strongly conserved, which is reflected in a higher mean MI for the MS2[S37P] than MS2[WT] (−0.24 and −0.33, respectively). This may seem counterintuitive, given that the mean AFS of the MS2[S37P] library was lower than that of the MS2[WT] library (−1.14 and −0.92, respectively). However, the lower mean AFS of the MS2[S37P] library results from the abundance of mutants at the minimum −4.0 AFS.

Disaggregation of AFS distribution by structural context (core, interface, or surface) reveals that the capsid AFLs differ most at subunit interfaces (Figure 5a). The MS2[WT] capsid shows a hierarchy whereby core residues are most conserved, followed by subunit interface residues, with surface residues (at either the interior or exterior surface) being most permissive to mutation. These preferences are in line with previous trends determined from multiple sequence alignment of T=3 VLPs. ^33^ MS2[S37P] capsids also possess a highly conserved core; however the mutability indicies of the interface and surface residues are statistically similar. Closer examination of contact interface context reveals that interdimer contact residues have similar mutabilities in both MS2[WT] and MS2[S37P]. Meanwhile, residues along intradimer contacts or residues that participate in multiple interface contacts are more permissive to mutation in the MS2[S37P] capsid library (Figure 5b).

**Figure 5:**
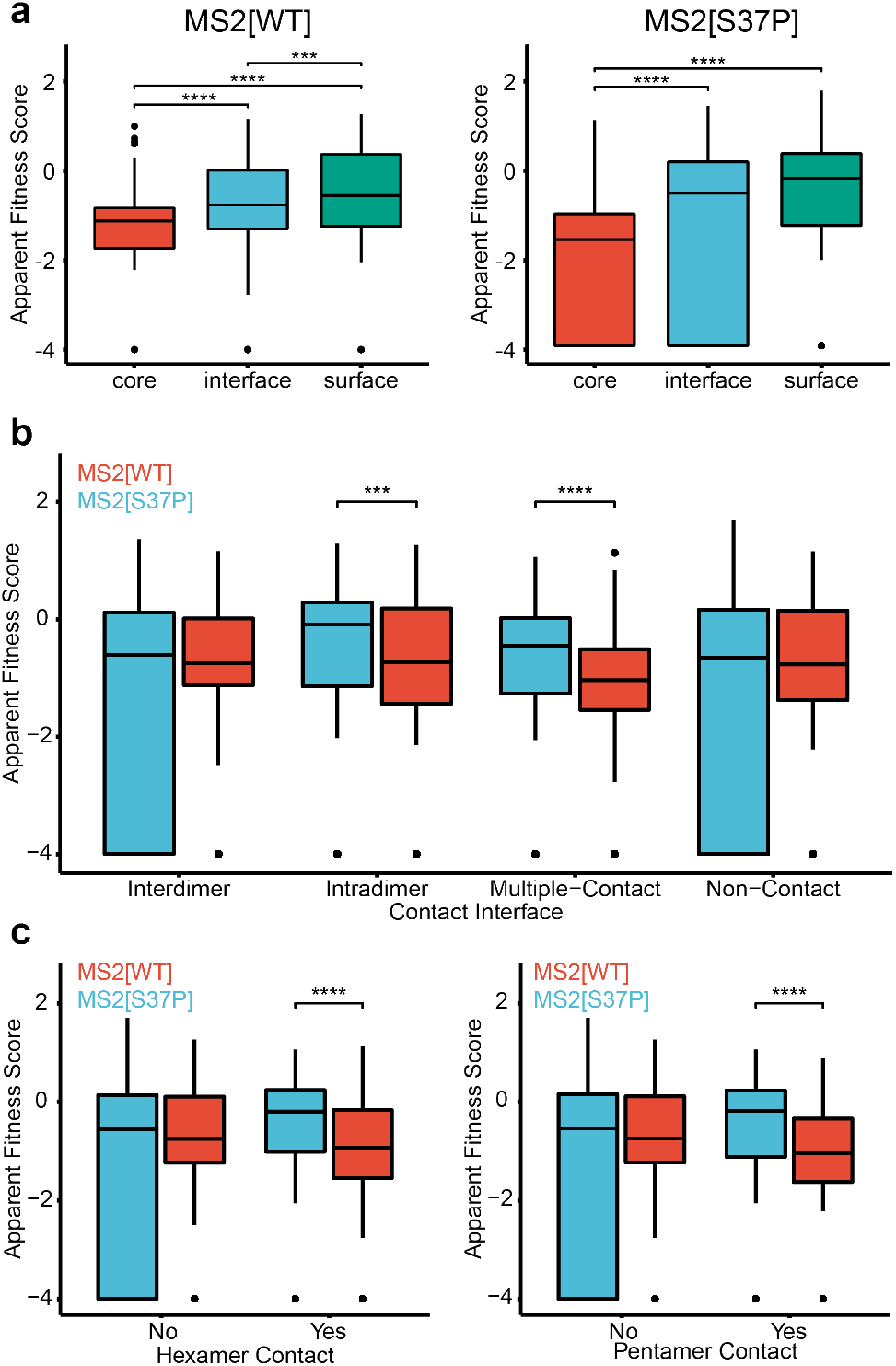
Capsid mutability trends based on structural context. (a) Boxplots of MS2[WT] and MS2[S37P] apparent fitness score (AFS) distributions by residue context. (b) Boxplot of AFS distributions by interface context, further separated in (c) by symmetry contact. MS2[WT] and MS2[S37P] distributions are shown in red and blue, respectively. *** indicates *p* < 0.001 and **** indicates *p* < 0.0001.

The symmetry elements of MS2[WT] and MS2[S37P] were next explored to understand better differences in the AFL of both capsid phenotypes. Casper and Klug’s theory of quasi-equivalence dictates that T=3 capsids will form from a mixture of pentameric and hexameric subunits at 5-fold and quasi-6-fold axes of symmetry, while T=1 icosahedral capsids form only from pentamer subunits at 5-fold symmetry axes.^34^ Prior work has also isolated two key intermediates in the MS2[WT] assembly pathway: a 12mer subunit at the quasi-6-fold symmetry axis (6CP2) and a 20mer subunit at the 5-fold symmetry axis (10CP2).^35^ While assembly intermediates of MS2[S37P] have not been directly observed, the 10CP2 subunit is a viable assembly intermediate, while the 6CP2 subunit presumably is not due to the geometry of the T=1 capsid. As such, we hypothesized that residues involved in the pentameric interfaces of the MS2 capsid would be similarly conserved in MS2[WT] and MS2[S37P], while the hexameric interfaces would be more flexible in MS2[S37P] as they do not mediate productive intermediates in the assembly pathway. Surprisingly, we find that both hexameric and pentameric interface residues are significantly more flexible in MS2[S37P]. This may suggest that the assembly pathway of the T=1 capsid may be more reliable than the T=3 capsid once initial folding and dimerization has occurred. There are indeed more potential misassembly pathways in T=3 icosehedra due to their larger number of unique interfaces, ^33^ although the relationship between the number of improper assembly pathways and the mutational flexibility of VLPs has not been previously established. These trends and observations demonstrate how comprehensive fitness landscapes may reveal mutability preferences that challenge expectations from sequence conservation in nature or known assembly pathways.

### Benchmarking Computational Residue Scanning Methods for Variant Fitness Prediction

We used these datasets to benchmark computational tools that calculate predicted changes in folding free energy of protein variants. *In silico* deep mutational scans of the MS2[WT] and MS2[S37P] coat proteins were performed using BioLuminate,^36^ the DynaMut2 web server^37^ and the PoPMuSiC 2.1 web server.^38^ As modeling the monomer subunit only accounts for one potential mode of assembly failure (protein misfolding), computational scans of the dimeric, 6CP2, and 10CP2 subunits of the assembled capsid were performed to achieve better coverage of the assembly deficits related to changes in intra-dimer and inter-dimer affinity. The 6CP2 capsomer was not modeled for MS2[S37P] as it is not likely to be formed in the assembly pathway. Correlation between computationally-predicted ΔΔG of coat protein variants and experimentally determined AFS was low across all tested methods and modeled subunits. This suggests that calculated fitness values of the capsid assemblies depends on a more complex set of factors than are captured by the endpoint Gibbs free energy of its constituent subunits.

Although the absolute AFS is not well correlated with computational ΔΔG scoring, we hypothesized that these methods may be successful in a binary classification of variant assembly competency. Previously-established AFS thresholds for assembly were used to label variant phenotypes (AFS > 0.2 = “assembling”, AFS < –0.2 = “non-assembling”). Predictions were classified as “assembling” if ΔΔG <= 0 (i.e. computation predicts the mutation is neutral or stabilizing) and “non-assembling” if ΔΔG > 0 (i.e. computation predicts the mutant is destabilizing). Binary classification results are shown in Figure 6. The accuracy of the tested classification methods was modest, ranging from 0.62-0.73. There were no large differences in accuracy between the three computational scanning methods. Gratifyingly, models of assembly subunits improved classification accuracy over models of the coat protein monomer for both MS2[WT] and MS2[S37P] in all three computational methods. Accuracy can, however, provide an overly-optimistic measure of performance in cases where the tested dataset has a class imbalance (i.e. there are many more non-assembling mutations than assembling mutations). ^39^ Thus, the Matthews correlation coefficient (MCC) was reported for each classification case, as this metric is a more reliable measure for classification of imbalanced datasets.^40,41^ MCC was calculated with values derived from Table 1 using equation 1. MCC ranges from −1 to +1, where MCC = +1 represents a perfect classifier, MCC = −1 represents perfect misclassification, and MCC = 0 represents the expected value of a coin toss classification.

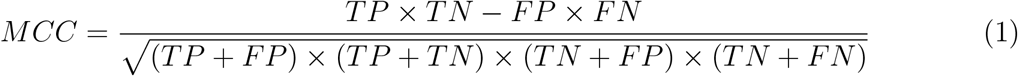

**Figure 6:**
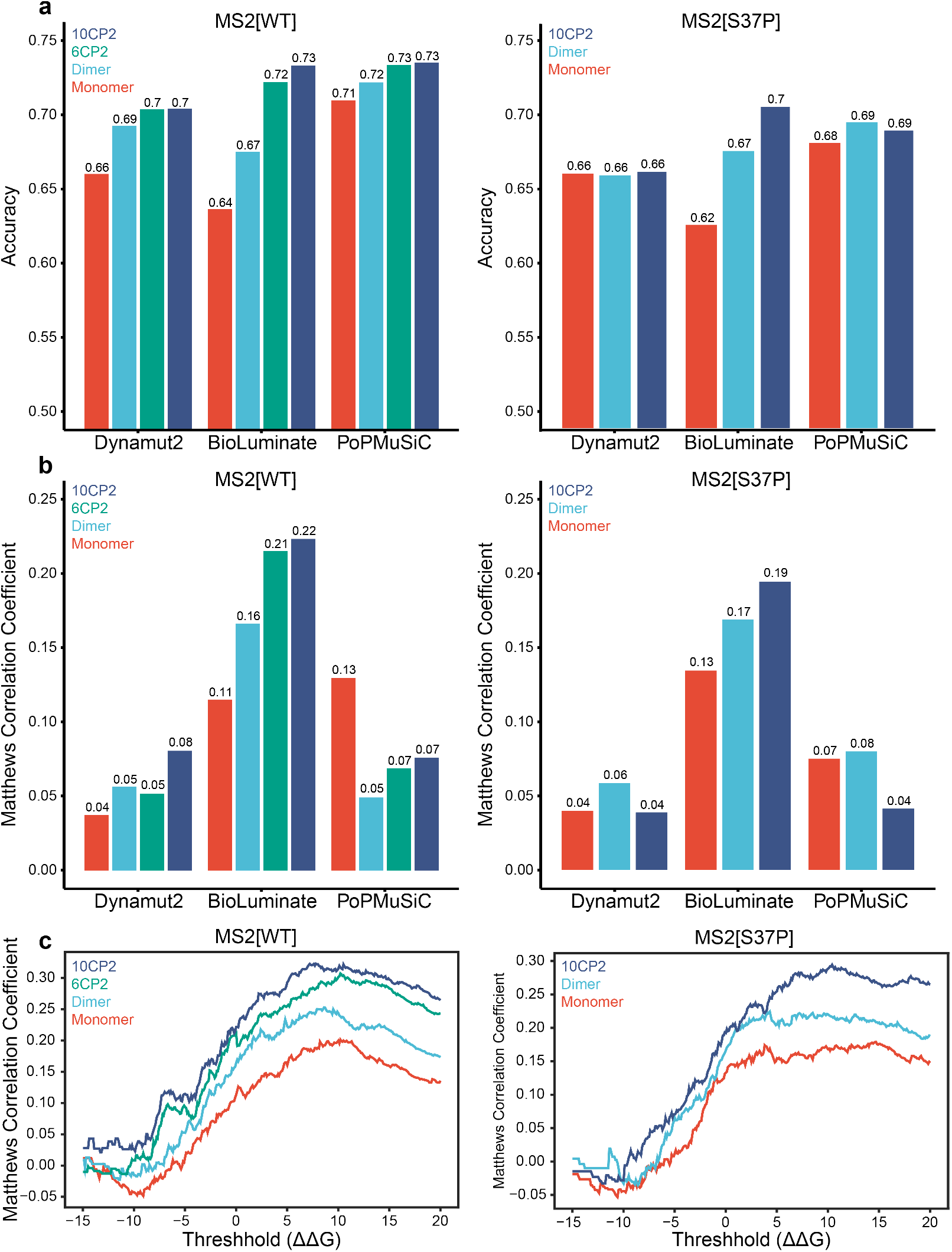
Computational residue scanning classification performance. The (a) accuracies and (b) Matthews correlation coefficients of three computational residue scanning methods are shown for various subunit models of the MS2[WT] and MS2[S37P] capsid. Predicted ΔΔG values from BioLuminate’s ΔStability parameter were compared to experimental apparent fitness scores. (c) Matthews correlation coefficients are shown as a function of classification ΔΔG threshhold.

**Table 1:** Confusion matrix describing performance of computational scanning classification.

|  |  | Fitness Landscape |  |
| --- | --- | --- | --- |
|  |  | Assembling | Non-Assembling |
| Computational Scan | Assembling | $TP$ | $FP$ |
| | Non-Assembling | $FN$ | $TN$ |

Evaluation by MCC reveals that the physics-based prediction via BioLuminate scored much higher than the other two methods (Figure 6b. While the overall accuracy of each method was relatively similar, BioLuminate correctly predicted the minority class (i.e., “assembling” mutation) in more cases than DynaMut2 and PoPMuSiC 2.1. Subsequent tuning efforts focused on using the predicted ΔStability from BioLuminate, since predicting assembly-competent mutations is more valuable for the production of capsids with engineered properties or new sequence motifs. In order to improve classifier performance, the ΔΔG threshold for designation of a mutation as “assembling” or “non-assembling” was scanned to optimize the MCC scores (Figure 6c). Threshold scanning revealed that a higher ΔΔG cutoff (ΔΔG = +5-8 kcal/mol) resulted in a higher MCC score for capsid phenotype classification. This indicates that MS2 capsid assembly can readily occur in spite of moderate increases to the folding free energy of the capsid subunits. Though tuning the cutoff threshold improved model performance, we sought to improve predictive power further by employing machine learning to capitalize on the rich feature set derived from these computational methods.

### Development of Supervised Learning Models for Capsid Assembly Classification

A total of 106 features for the MS2[WT] set and 81 features for the MS2[S37P] set were prepared for each amino acid variant represented in the experimental fitness landscapes of each capsid. The description of each feature is provided in the Supporting Information. Initial models were prepared via four supervised learning algorithms (K-Nearest Neighbor, ^42^ logistic regression, ^43^ random forest,^44^ and XGBoost^45^) These models were trained with default hyperparameters using a stratified 75/25 train/test split of the landscape data. Models were evaluated by 10-fold cross validation, and binary classifier performance was measured using the MCC scores (Figure 7). The random forest and XGBoost models outperformed the K-nearest neighbor and logistic regression models in both training sets, as measured by MCC. Bayesian Optimization was subsequently implemented to tune the hyperparameters used for XGBoost or random forest model training. ^46^ The resultant tuned models showed strong predictive performance on the unseen test set of landscape data, with the MS2[WT] model slightly outperforming the MS2[S37P] model (accuracy = 0.86, MCC = 0.58 and accuracy = 0.82, MCC = 0.52, respectively).

**Figure 7:**
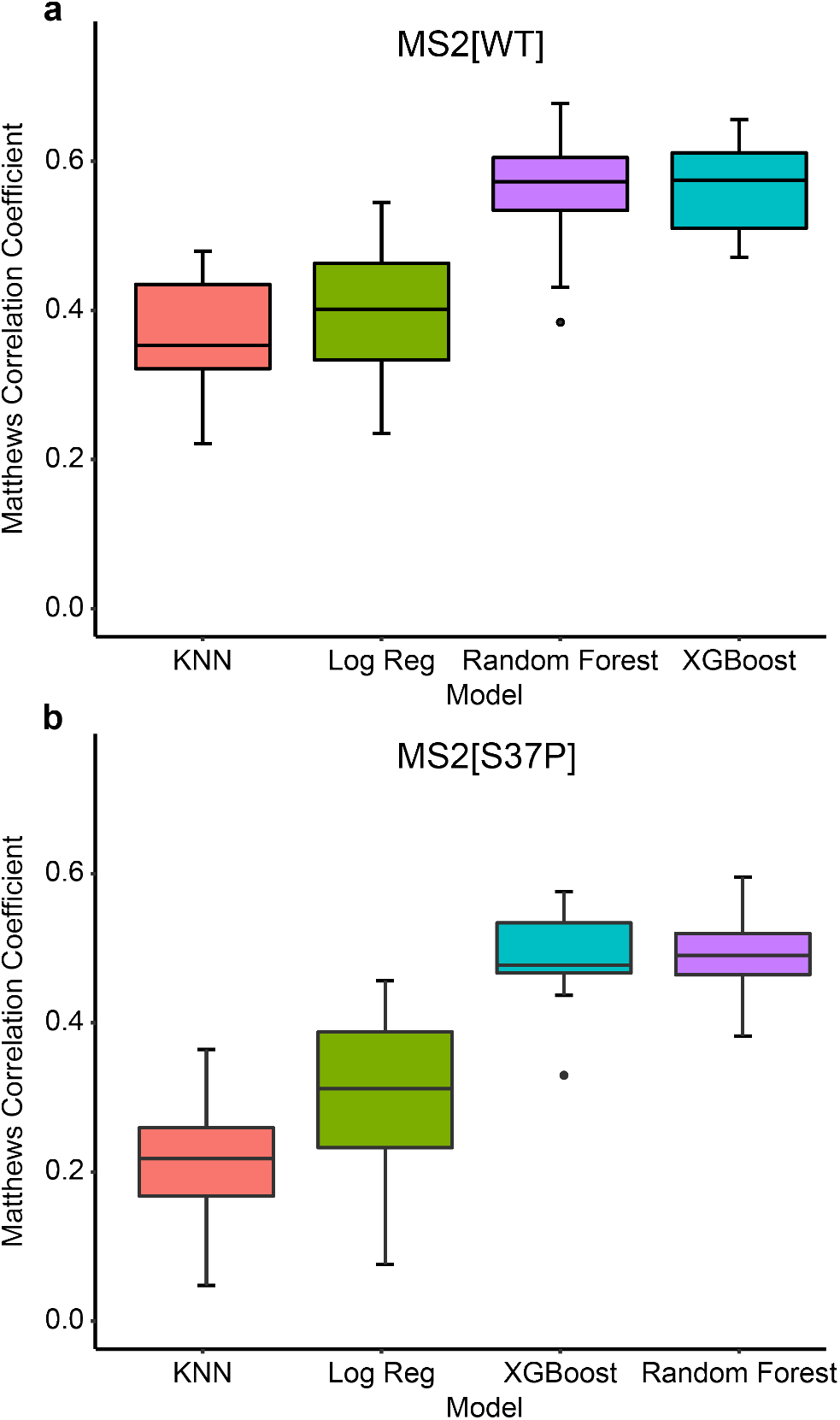
Initial machine learning model classification performance. Four machine learning algorithms were benchmarked for predicting the assembly state of (a) MS2[WT] or (b) MS2[S37P] capsid variants. Boxplots of the resultant Matthews correlation coefficients from 10-fold cross validation of models are shown.

Ensemble tree-based algorithms, such as XGBoost and random forest, furthermore enable calculation of the importance of each feature with respect to prediction of the target variable, thus offering interpretability of the final MS2[WT] and MS2[S37P] models (Figure 8).^47,48^ Across all four models, the most important feature for prediction was the ΔStability of the 10CP2 subunit calculated using Bioluminate. Many *de novo* features, such as residue solvent-accessible surface area, hydropathy, potential energy, and internal energy, contributed significantly to the final models. In the MS2[WT] models, ΔStability of the 6CP2 subunit was much less important than the 10CP2 subunit, which may indicate either that the endpoint stability of the 6CP2 complex does not capture inhibition of the subunit assembly pathway, or that this complex is more resilient to perturbation than the 10CP2 complex. Interestingly, no intersubunit affinity terms ranked highly in importance, while intradimer affinity ranked in the top ten of 2 out of 4 of the final models.

**Figure 8:**
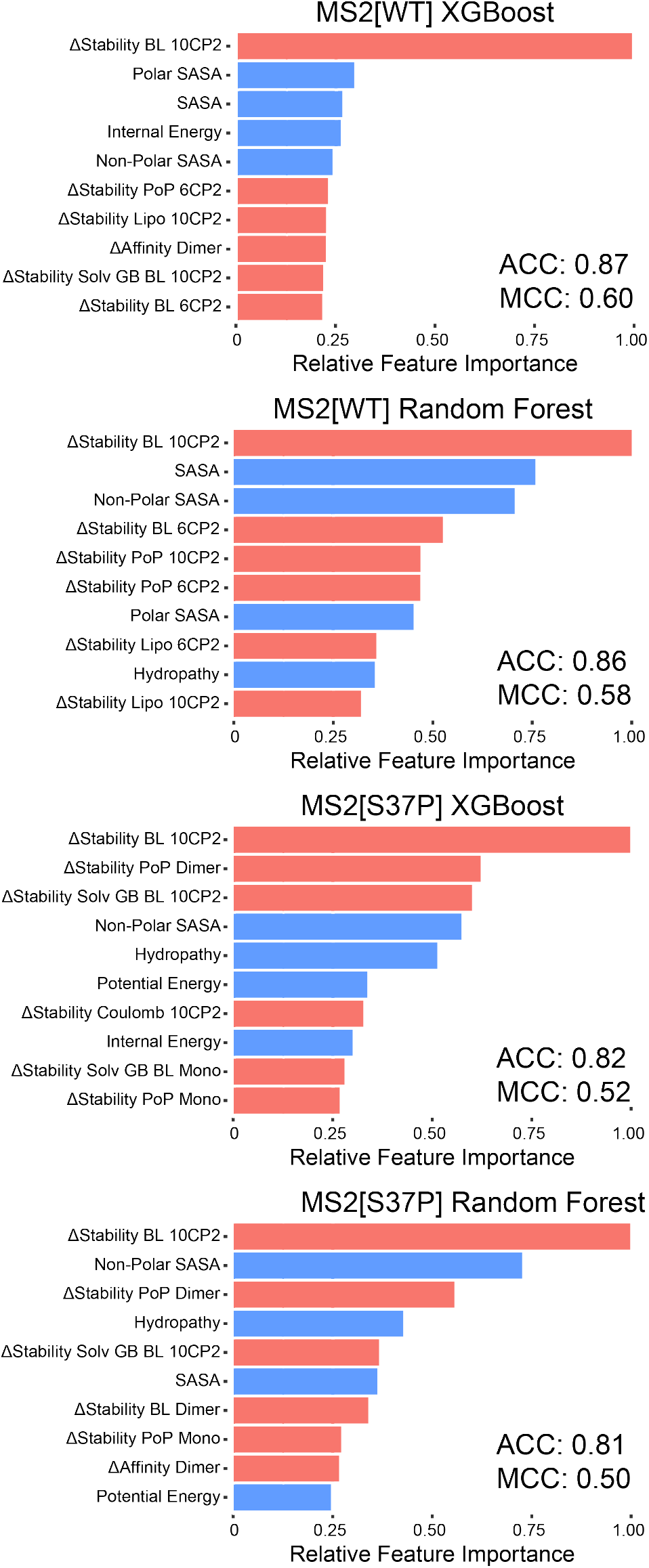
Final model performance and feature importance. Performance of XGBoost and random forest models of (a,b) MS2[WT] and (c,d) MS2[S37P] assembly classifications are shown. The top 10 most informative features for classification are shown. Model-derived and *de novo* features are shown in red and blue, respectively.

We next sought to establish the minimal input data set necessary for model training, as reduction of the training set size or initial computational modeling steps may enable accurate construction of assembled capsid fitness landscapes with reduced experimental and computational cost. Training features were reduced to include only *de novo* amino acid features as well as the most predictive feature, ΔStability derived from the 10CP2 Bioluminate modeling. A random forest model was trained on randomly-generated subsets of the fitness landscape (1-2000 training samples) and evaluated by predicting the remaining landscape assembly states. Learning curves of both models displaying the gain in MCC and accuracy as a function of training samples are shown in Figure 9. Interestingly, the model achieves reasonable accuracy after very few samples, and only marginally improves as the sample numbers increase. Viewing MCC gain, however, displays a sharp improvement that plateaus after roughly 500 samples. This suggests that small sublibraries of self-assembled protein capsids may enable reasonably accurate predictions of capsid fitness landscapes, likely by learning that the majority of mutations inhibit capsid assembly. To obtain predictions that correctly classify assembly-competent mutants, a moderate sampling of the full mutational space of the VLP (about 20% of variants) will be needed.

**Figure 9:**
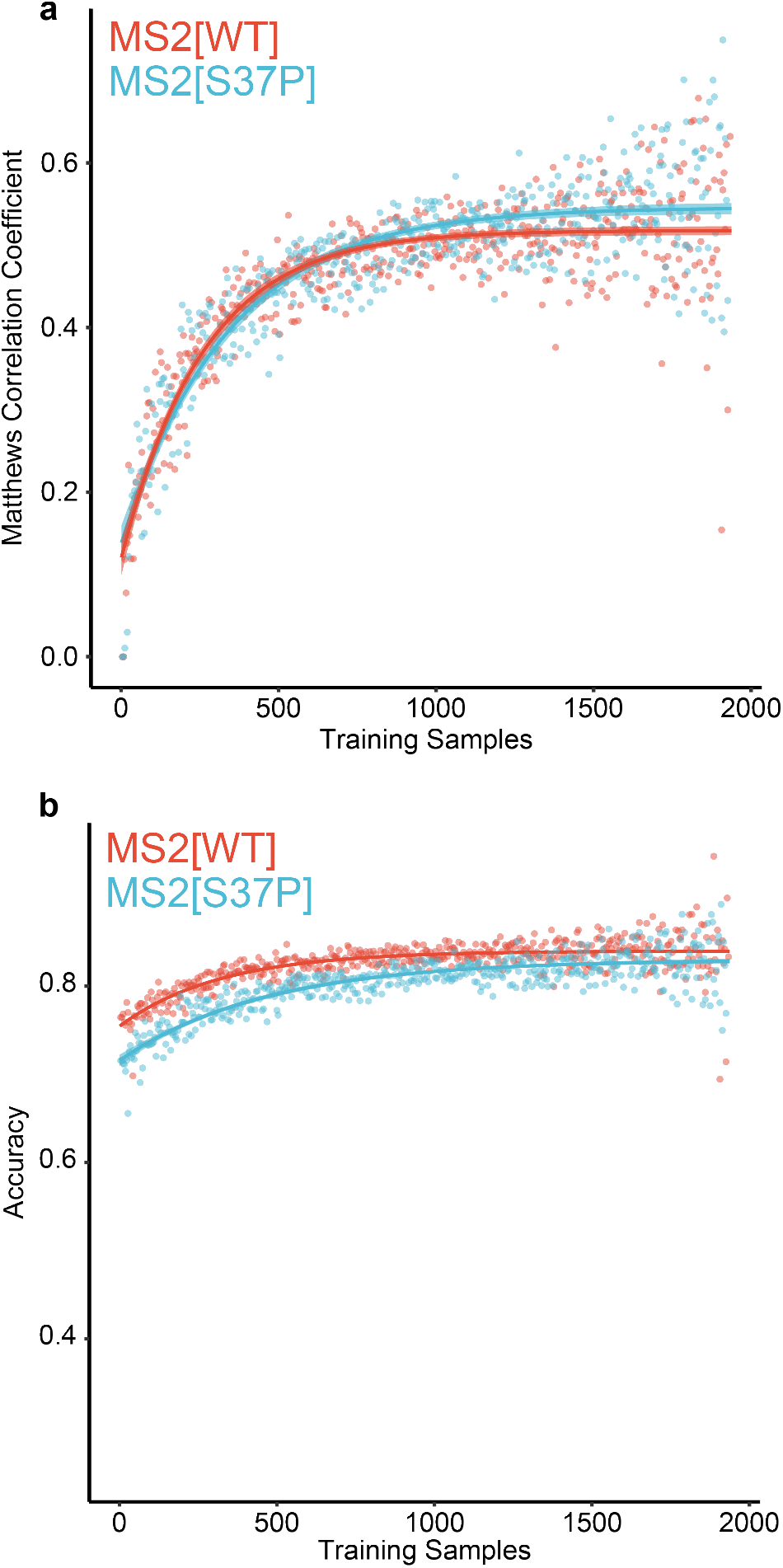
Determination of minimized input model performance. (a) MCC gain plot and (b) accuracy gain plot of both capsid models as functions of training set size. Models were generated using only ΔStability BL 10CP2 and *de novo* parameters for training. Random subsets of the fitness landscape data were used to simulate low-sample training information. MS2[WT] and MS2[S37P] learning curves are shown in red and blue, respectively.

## Conclusion

Here, we have characterized the apparent fitness landscape of a non-native (T=1 icosahedral) VLP assembly geometry. Comparison to the apparent fitness landscape of the wild type (T=3 icosahedral) particle yielded insight into the influences quaternary structure has on the mutability of a self-assembling material. Residues at the core of the monomer subunit, as well as residues mediating the dimer interfaces of the capsid, showed similar mutability preferences in both capsid libraries. Surprisingly, residues along both the five-fold and quasi-six-fold symmetry interfaces of the particle were significantly more mutable in the T=1 capsid than the wildtype assembly, suggesting that the T=1 capsid may be a more versatile platform for engineering efforts. Further, the deep mutational scanning data produced were used to benchmark the performance of various computational residue scanning tools for estimating the effects of missense mutations on protein folding energetics. While correlations between calculated ΔΔG values and measured assembly competency were relatively low, modeling known assembly intermediates using BioLuminate’s MM/GBSA method offered the best overall performance. Finally, machine learning models were trained using *de novo* structural features in tandem with computational modeling data to produce predictive assembly state classifiers for both capsid assemblies. The relationship between input training size and classifier performance demonstrates that relatively small input datasets can be used to produce predictive classifiers for self-assembly with strong performance. As such, this study may represent a generalizable strategy for semi-rational engineering and systematic mutation of proteinaceous self-assembled nanomaterials.

## Experimental

### Strains

All library experiments were conducted with MegaX DH10B *E. coli* electrocompetent cells (ThermoFisher Scientific, Cat# C640003). Overnight growths from a single colony were incubated for 18 h at 37 °C with shaking at 200 RPM in LB-Miller media (Fisher Scientific, Cat# BP1426-2) with chloramphenicol at 32 mg/L. Expressions were subcultured 1:100 into 2xYT media (Teknova, Cat# Y0210) with 32 mg/L chloramphenicol, allowed to grow to an OD600 of 0.6, then induced with 0.1% arabinose. Expressions continued overnight at 37 °C with shaking at 200 RPM.

### FPLC SEC

MS2 CP libraries and select individual variants were purified on an Akta Pure 25 L Fast Protein Liquid Chromatography (FPLC) system with a HiPrep Sephacryl S-500 HR column (GE Healthcare Life Sciences, Cat# 28935607) Size Exclusion Chromatography (SEC) column via isocratic flow with 10 mM phosphate buffer at pH 7.2 with 200 mM sodium chloride and 2 mM sodium azide.

### HPLC SEC

Variants or libraries were analyzed on an Agilent 1290 Infinity HPLC with an Agilent Bio SEC-5 column (5 um, 2000A, 7.8×300mm) with isocratic flow of 10 mM phosphate buffer at pH 7.2 with 200 mM sodium chloride and 2 mM sodium azide. Fractions were harvested at 11.2 min, or the characteristic elution time for wild-type MS2. Harvested VLPs were then subjected to RNA extraction and high-throughput sequencing sample preparation.

### Library generation

Deep mutational scanning libraries of both capsids were performed as previously described via SyMAPS.^23^ SyMAPS uses the EMPIRIC cloning developed in the Bolon lab.^49^ In EMPIRIC cloning, a plasmid contains a self-encoded removable fragment (SERF) flanked by inverted BsaI recognition sites. Thus, BsaI digestion simultaneously removes the SERF and BsaI sites. Five previously-generated entry vectors each designed to accept a 26 codon region of the MS2 coat protein as an insert were used to prepare both the MS2[WT] and MS2[S37P] libraries. Four of the entry vectors bearing the position S37 codon were mutated to S37P via Quikchange mutagenesis for the MS2[S37P] library. Single-stranded DNA primers spanning each 26-codon region with an NNK codon at each position were pooled and diluted to a final concentration of 50 ng/uL. Touchdown PCR was used to amplify the mutagenic inserts and the double-stranded DNA was purified by PCR clean-up (Promega, Cat# A9282). Cleaned dsDNA inserts were diluted to 5 ng/μL, then cloned into the entry vector using Golden Gate cloning. Ligated plasmids were incubated on desalting membranes (Millipore Sigma, Cat# VSWP02500) for 20 min, followed by transformation into electrocompetent DH10B E. coli cells (Invitrogen, Cat# 18290015). Following electroporation and recovery, cells were plated onto LB-A containing 32 μg/mL chloramphenicol and grown overnight at 37 °C. Dilutions (100-fold) of each replicate were plated and counted to ensure a minimum of 3x colony-forming units relative to the number of possible variants.

### Assembly selection

Colonies were harvested by scraping plates into LB-M and growing the mixture for 2 h. Each library was subcultured 1:100 into 1 L of 2xYT (Teknova, Cat# Y0210) and grown to an OD of 0.6, then induced with 0.1% arabinose. Variant libraries were expressed overnight at 37 °C. Cells were harvested by centrifugation and lysed by sonication. Libraries were subjected to two rounds of ammonium sulfate precipitation (50% saturation), followed by FPLC size exclusion chromatography purification to select for well-assembled T=3 or T=1 VLPs. Collected fractions were then subjected to 50 °C thermostability challenges in a H_2_O bath for 10 min. Precipitated VLPs were pelleted via centrifugation, and assembled VLPs were isolated via semi-preparative HPLC size exclusion chromatography. Fractions containing assembled VLPs (at the characteristic elution time of 6.5 min for MS2[WT] or 8.0 min for MS2[S37P]) were combined, subjected to RNA extraction, barcoded, and identified via high-throughput sequencing.

### High-throughput sequencing sample preparation

Plasmid DNA was isolated prior to expressions using Zyppy Plasmid Miniprep Kit (Zymo, Cat# D4036). RNA was extracted from the assembly-selected libraries using previously-published protocols. ^50^ Briefly, homogenization was carried out with TRIzol (Thermo Fisher Cat# 15596026), followed by chloroform addition. The sample was centrifuged to separate into aqueous, interphase, and organic layers. The aqueous layer, containing RNA, was isolated, and the RNA was precipitated with isopropanol and washed with 70% ethanol. RNA was dried and resuspended in RNAse free water. cDNA was synthesized with the Superscript III first strand cDNA synthesis kit from Life (Cat# 18080051, random hexamers primer).

cDNA and plasmids were both barcoded for high-throughput sequencing. Both types of samples were amplified with two rounds of PCR (10 cycles, followed by 8 cycles) to add barcodes and Illumina sequencing handles, following Illumina 16S Metagenomic Sequencing Library Preparation recommendations. Libraries were quantified with Qubit and combined in equal molar ratio. Libraries were analyzed by PE300 MiSeq in collaboration with the UC Davis Sequencing Facilities.

### High-throughput sequencing data processing and analysis

Data were trimmed and processed described previously, ^23^ with minor variation. Briefly, data were trimmed with Trimmomatic^51^ with a 4-unit sliding quality window of 20 and a minimum length of 32. Reads were merged with FLASH (fast length adjustment of short reads) with a maximum overlap of 300 base pairs. Reads were sorted and indexed with SAMtools^52^ and unmapped reads were filtered with the Picard function CleanSam. Reads were processed with in-house code to produce various AFLs.

### AFL calculations

Cleaned and filtered high-throughput sequencing reads were analyzed using in-house Python code (available at https://github.com/dan-brauer/MS2-Assembly-Classification). Briefly, the start codon and stop codons on either end of the MS2 gene were parsed and the correct gene length was verified. For MS2[S37P], codon encoding for the S37P mutation was also verified. Codons were then translated into amino acids to generate counts for each tripeptide combination

Calculations were repeated for all replicate plasmid pools and assembled selections. Relative abundances were calculated similarly to the previously described protocol.^26^ Briefly, the relative abundance of a selection was calculated by dividing the abundance generated from an assembled VLP by the plasmid abundance from its respective biological replicate.

The mean relative abundance across three biological replicates was calculated for the final fitness landscape of each library. All NaN (null) values, which indicate variants that were not identified in the plasmid library were ignored. Scores of zero, which indicate variants that were sequenced in the unselected library but absent in the VLP library, were replaced with an arbitrary score of 0.0001. The log10 relative abundance value for an individual variant in a particular challenge is referred to as its Apparent Fitness Score (AFS).

### Computational fitness scanning

Assembly subunits (monomer, dimer, 6CP2, and 10CP2) were constructed in the Maestro molecular modeling environment based on the crystal structures of MS2[WT] (PDB:2MS2) and MS2[S37P] (PDB:4ZOR). Resultant structures were preprocessed and minimized using the OPLS3e forcefield. Computational scanning for minimized subunit structures via DynaMut2 and PoPMuSiC 2.1 was performed using their respective web servers. BioLuminate calculations were performed symmetrically across chains using side-chain prediction with backbone minimization at a 0.00 A cutoff. Classifier performance was evaluated using in-house Python code (available at https://github.com/dan-brauer/MS2-Assembly-Classification).

### Machine learning model training

Machine learning models were trained and assessed using the Tidymodels ^53^ framework in R (available at https://github.com/dan-brauer/MS2-Assembly-Classification). Continuous training variables were normalized, nominal variables (e.g. amino acid identity) were encoded as binary dummy variables, and zero variance variables were filtered in data preprocessing. Algorithm comparisons (K-Nearest Neighbor, logistic regression, random forest, and XG-Boost) were performed using default hyperparameters. Initial random forest and XGBoost model training used 1000 trees. Cross validation (10-fold) was performed on a 75/25 split of the fitness landscaping data. Random forest and XGBoost models were further optimized through 100 rounds of Bayesian hyperparameter tuning, with 10 initial parameter sets and an early stop after 20 consecutive iterations with no improvement. Optimized parameters were used to train final models which were evaluated on the 25% of data held out for final evaluation. Model feature importance was determined using the R ‘vip’ package.^54^

## Supporting information

Supplemental Data Wildtype

## Acknowledgement

This work was funded in part by the National Science Foundation (grants CBET-2043973 to DTE and CBET-2044011 to MBF). The authors thank Dr. Kathleen Durkin at the UC Berkeley Molecular Graphics and Computation Facility for assistance in running BioLuminate computational Scanning through Maestro. The sequencing was carried out at the DNA Technologies and Expression Analysis Cores at the UC Davis Genome Center, supported by NIH Shared Instrumentation Grant 1S10OD010786-01.

